# Extracellular monomeric and aggregated tau efficiently enter human neurons through overlapping but distinct pathways

**DOI:** 10.1101/168294

**Authors:** T. Wassmer, L. Evans, G. Fraser, J. Smith, M. Perkinton, A. Billinton, F.J. Livesey

**Author notes:** Co-first authors (T. Wassmer, L. Evans).

## Abstract

A current working model is that Alzheimer's disease actively spreads from diseased to healthy neurons, mediated by transfer of extracellular, abnormal, disease-specific forms of the microtubule-associated protein tau. It is currently unclear whether transfer of tau between neurons is a toxic gain-of-function process in dementia, or reflects a constitutive biological process. We report two mechanisms of entry of monomeric tau to neurons: a rapid early dynamin-dependent phase, and a second, slower actin-dependent phase, suggesting that monomeric tau enters neurons via rapid saturable clathrin-mediated endocytosis and also by bulk endocytosis. Aggregated tau entry is independent of actin polymerisation and largely dynamin dependent, consistent with clathrin-mediated endocytosis and distinct from macropinocytosis, the major route for aggregated tau entry reported for non-neuronal cells. Anti-tau antibodies abrogate tau entry into neurons, with tau carrying antibody with it into neurons, indicating that antibody binding is insufficient to prevent neuronal tau entry.

## Introduction

The microtubule-associated protein tau (MAPT) is involved in the pathogenesis of several forms of dementia, including Alzheimer’s disease (AD) and frontotemporal dementia (FTD). Mutations in the MAPT gene are causal for some familial forms of FTD (Ghetti et al., 2015), and the formation of intracellular, hyperphoshorylated aggregates of tau (neurofibrillary tangles, NFTs) is a common pathological feature in AD, FTD and other dementias (Grundke-Iqbal et al., 1986; Kosik et al., 1986). Disease progression, and clinical severity of symptoms, is associated with a predictable spatial and temporal order of appearance of NFTs in different forebrain regions (Braak and Braak, 1991). This has led to the proposal that these diseases actively spread from diseased to healthy neurons in a spatial and temporal progression, mediated by extracellular, abnormal, disease-associated forms of tau (Iba et al., 2013; Liu et al., 2012).

It is not currently known if neuronal release and internalisation of extracellular tau are disease-specific phenomena that spread disease through the CNS, or are fundamental physiological processes. *In vivo*, tau protein is present in interstitial fluid of the central nervous system (CNS) and passes into cerebrospinal fluid (CSF), where is it is found in concentrations of the order of 15pM (Blennow et al., 1995; Olsson et al., 2005). Mice transgenic for human tau have 10-fold higher concentrations of tau in brain interstitial fluid than in CSF, which would suggest that the extracellular concentrations of tau in the human brain are in the high picomolar-low nanomolar range (Yamada et al., 2011). There is strong evidence for regulated release of non-pathogenic forms tau from healthy neurons, and of tau entry into neurons and non-neuronal cells (Bright et al., 2015; Chai et al., 2012; Kanmert et al., 2015; Wang et al., 2017). There have been conflicting reports about the ability of extracellular monomeric tau to enter cells, whereas aggregated or fibrillar tau has been clearly shown to efficiently enter neurons and other cell types (Frost et al., 2009; Kfoury et al., 2012; Wu et al., 2013).

We used human stem cell-derived neurons to address open questions about the efficiency with which tau enters human neurons, which forms (monomeric and aggregated) of tau enter neurons, and the routes by which they do so. We find that both forms of tau are efficiently taken up by human neurons, by overlapping but distinct mechanisms, consistent with regulated endocytosis. Monomeric tau enters neurons by two different routes and is actively trafficked within the neuron. Tau entry into neurons can be slowed by antibody binding, however, extracellular tau-antibody complexes are internalised into intracellular compartments.

## RESULTS

### Rapid and efficient entry of extracellular monomeric and aggregated tau into human neurons

We first analysed the ability of monomeric or aggregated tau (P301S) protein to enter human cortical neurons from the extracellular milieu. We used the tau P301S variant, an autosomal dominant mutation that causes early onset FTD with high penetrance (Bugiani et al., 1999; Guo et al., 2017), enabling us to study the transmission of a disease-relevant variant that differs from normal tau by a single amino acid. To do so, purified recombinant monomeric (native) or heparin-aggregated tau protein conjugated to an amide reactive fluorophore (tau-Dylight 488 NHS ester) were incubated with iPSC-cell derived neurons for 2 hours. After extensive washing, monomeric and aggregated tau-Dylight were both detected in cells expressing the neuron-specific microtubule-associated protein MAP2, confirming that both forms of tau enter neurons (Figure 1A). Tau-Dylight was found predominantly within the somatic compartment of neurons (Figure 1A). After four hours incubation with extracellular tau, flow cytometry analysis (Figure 1B,C) revealed that 83% and 73% of dissociated cells contained monomeric or aggregated tau-Dylight, respectively, demonstrating that extracellular tau efficiently enters human neurons in culture.

**Figure 1.**
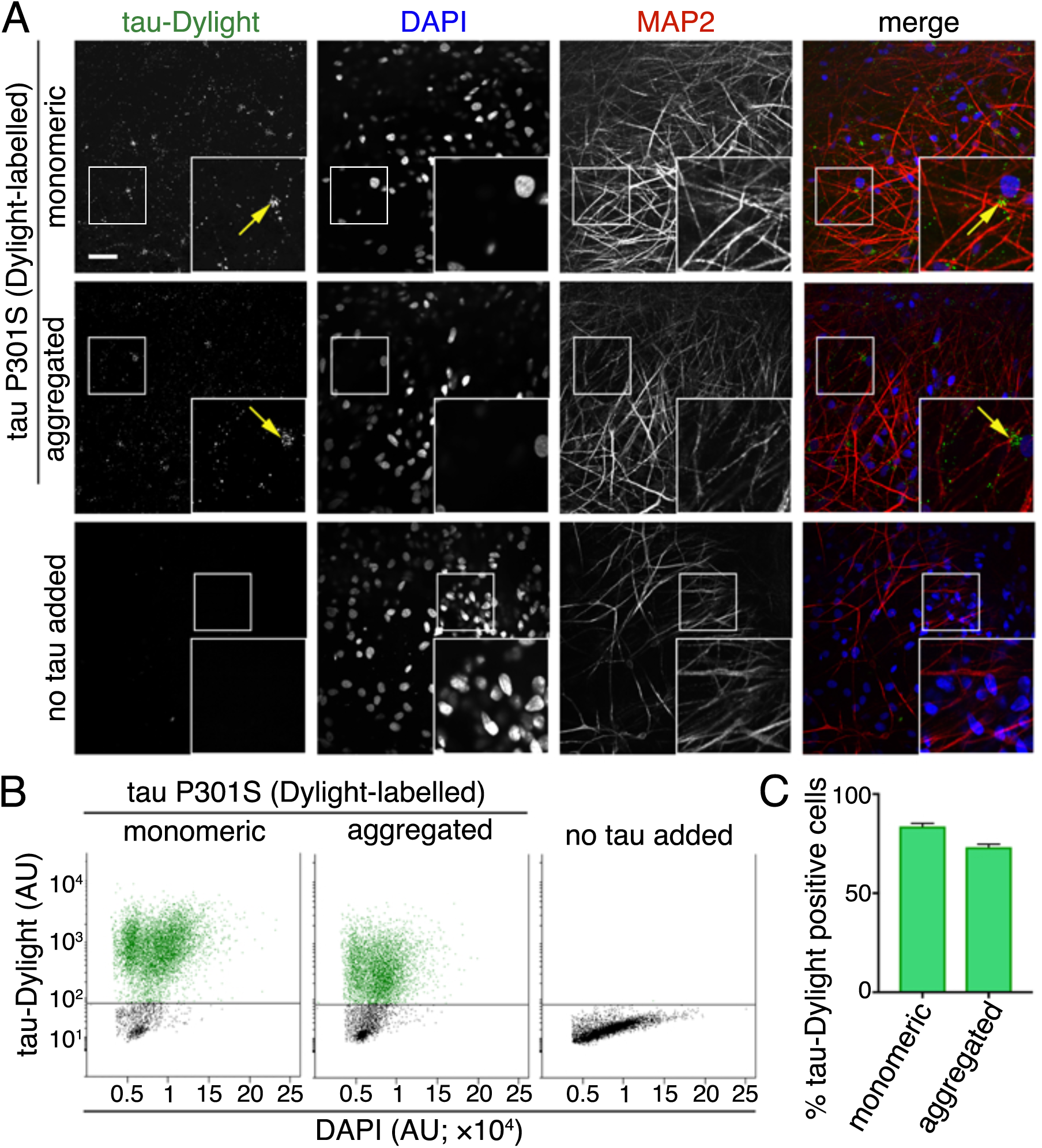
Human neurons efficiently take up extracellular monomeric and aggregated tau. (A) Immuno-fluorescent images showing neuronal uptake of extracellular monomeric or aggregated forms of tau P301S protein conjugated to an amide reactive fluorophore (green in merge; tau-Dylight) after 2 hours of incubation or without tau protein (no tau added). Fixed iPSC derived neurons (70 days after induction) were co-stained with nuclear and dendritic markers; DAPI (blue in merge) and MAP2 (red in merge), respectively. Fluorescent signals were captured using Opera Phenix system and representative images are shown. Scale bar 50μ. (B) Analysis of the proportion of tau positive neurons. iPSC neurons (63 days after induction) were incubated with monomeric or aggregated tau-Dylight protein for 4 hours before dissociation into single cells, counter staining with nuclear stain (DAPI) and analysis by flow cytometry. Scatterplots of DAPI /tau-Dylight double-labelled neurons were gated firstly by intensity of DAPI fluorescence and subsequently by tau-Dylight fluorescence, neurons without tau incubation (no tau added) were used to establish positive level tau-Dylight fluorescence. (C) Percentage of DAPI /tau-Dylight positive gated iPSC-derived neurons (total count of >7×10^3^ neurons) incubated with either monomeric, aggregated or without tau. Error bars indicate s.e.m, n=6.

### Concentration-dependent entry of tau into neurons via low pH transport vesicles

Live imaging of human cortical neurons was used to investigate the underlying mechanisms of neuronal entry of monomeric and aggregated tau. To do so, we prepared purified recombinant tau P301S protein conjugated to a maleimide reactive pH-sensitive dye (tau-pHrodo Red maleimide), a reporter detectable in low pH environments, including late endosomes and lysosomes. Incubation of tau-pHrodo with human neurons at a range of concentrations from 2.5 to 25 nM (0.12 to 1.2*μ*g.ml^−1^, diluted in culture medium) showed that tau entry to neurons is rapid and concentration dependent, as visualised by live imaging. Intraneuronal fluorescent punctae were observed within the first ten minutes of exposure to monomeric tau-pHrodo (Figure 2A, Supplementary movie 1). Tau-pHrodo positive structures increased in size and intensity over the four-hour course of the assay. These structures were present within neurites and accumulated in the somatic compartment of neurons. In the presence of 15 and 25 nM monomeric tau-pHrodo, the number of tau-pHrodo positive objects (Figure 2C) rapidly increases with the signal approaching a plateau (>90% of final measurement) after approximately one hour.

**Figure 2.**
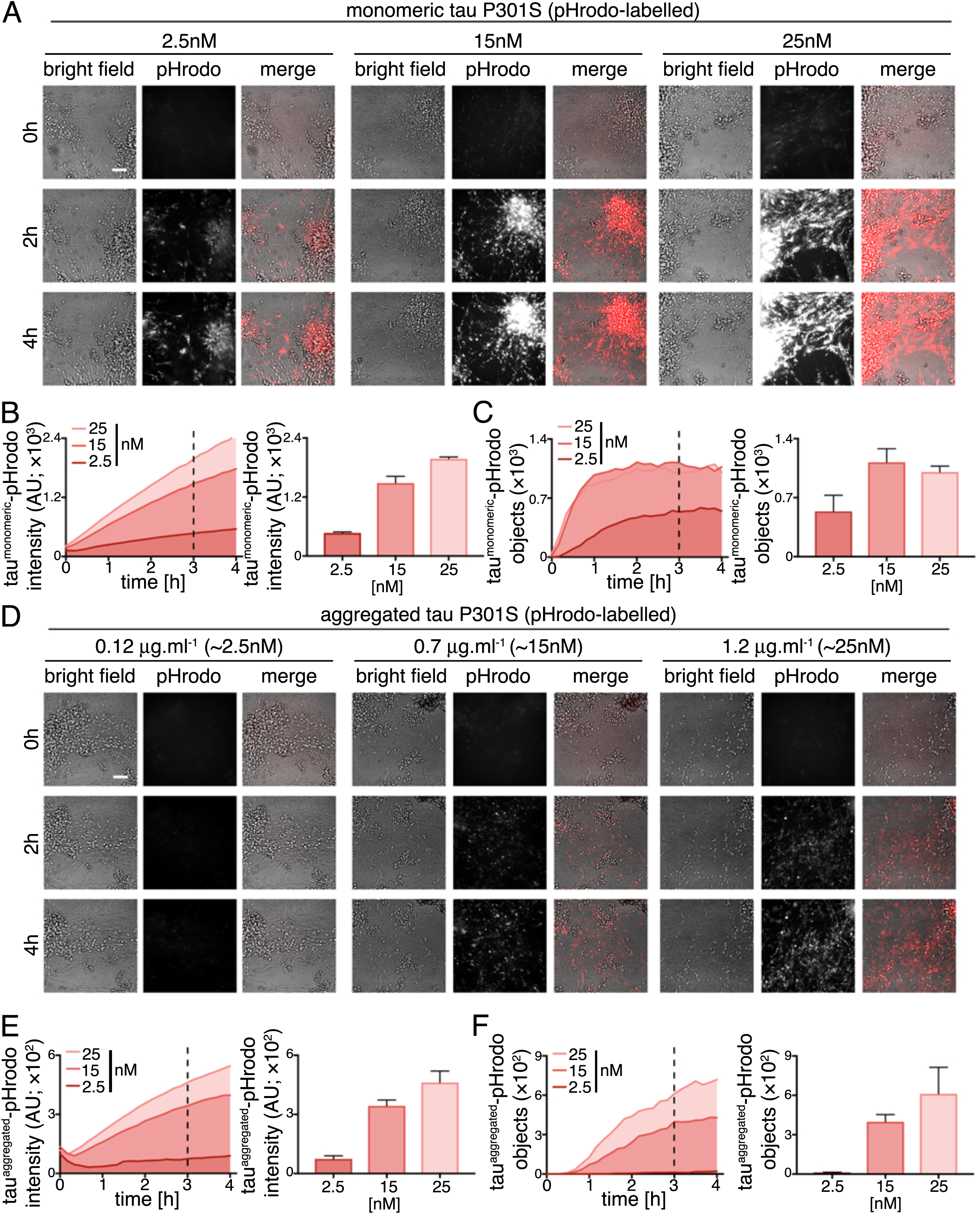
Human neurons take up extracellular tau via acidic intracellular compartments in a concentration-dependent manner. (A) Time-lapse (0-4 hours) images showing concentration dependent internalization and acidification of extracellular monomeric tau P301S conjugated to a pH-sensitive dye (tau-pHrodo [2.5, 15 or 25nM]; inverse relationship between fluorescence and pH) into iPSC derived human neurons (75 days after induction). Bright field (grey scale in merge) and pH sensitive fluorescent signal (pHrodo; red in merge) were captured using automated imaging on the Opera Phenix platform (Perkin Elmer). Nine independent measurements were taken from three technical replicates at 10 minutes intervals. Scale bar 100μ. After extracellular addition of 2.5, 15 or 25nM tau-pHrodo, (B) the sum intensity of the pHrodo positive objects and (C) the number of pHrodo positive objects was measured over 4 hours. Intensity (AU) and object measurements are displayed over time and at a three-hour time point (dashed line). Analysis was performed using Harmony software (Perkin Elmer) error bars indicate standard deviation. (D-F) Concentration dependent internalisation and acidification of 0.12, 0.7 or 1.2*μ*g.ml^−1^ of extracellular aggregated tau-pHrodo (equivalent to 2.5, 15 or 25nM of monomeric tau, respectively) into intraneuronal compartments using the same experimental conditions and parameters as (A-C).

Internalisation of aggregated tau-pHrodo (Figure 2D) was also found to be concentration dependent (Figure 2E). However, unlike monomeric tau-pHrodo, the number of detectable tau-pHrodo positive objects (Figure 2F) did not increase immediately, but displayed a lag for the first hour (Figure 2E). These kinetics of aggregated tau-pHrodo entry are similar to that of both lower concentrations of monomeric tau (2.5nM) and of low molecular weight (10kDa) dextran-pHrodo (same molarity as monomeric tau samples; Supplementary Figure 1). Direct quantitative comparison of the amount of monomeric and aggregated tau-pHrodo entering at different concentrations is not possible under these experimental conditions, as the mixed nature of aggregated tau fibrils prevents us establishing molarity accurately.

### Monomeric and aggregated tau appear in early endosomes and in lysosomes

To establish the route of vesicular internalisation of monomeric and aggregated tau, we analysed the appearance of tau-Dylight in endosomal or lysosomal compartments in iPSC-derived human neurons. Monomeric or aggregated tau-Dylight (total protein concentration 10μg.ml^−1^; ~250nM) was incubated with neurons for up to 6 hours, then extensively washed to remove extracellular protein and fixed. Co-localisation of tau with endosomes, defined by early endosomal antigen-1 (EEA1) immunofluorescence (Figure 3A), or late endosomes and lysosomes, defined by LAMP1 immunofluorescence (Figure 3B), was analysed by confocal microscopy.

**Figure 3.**
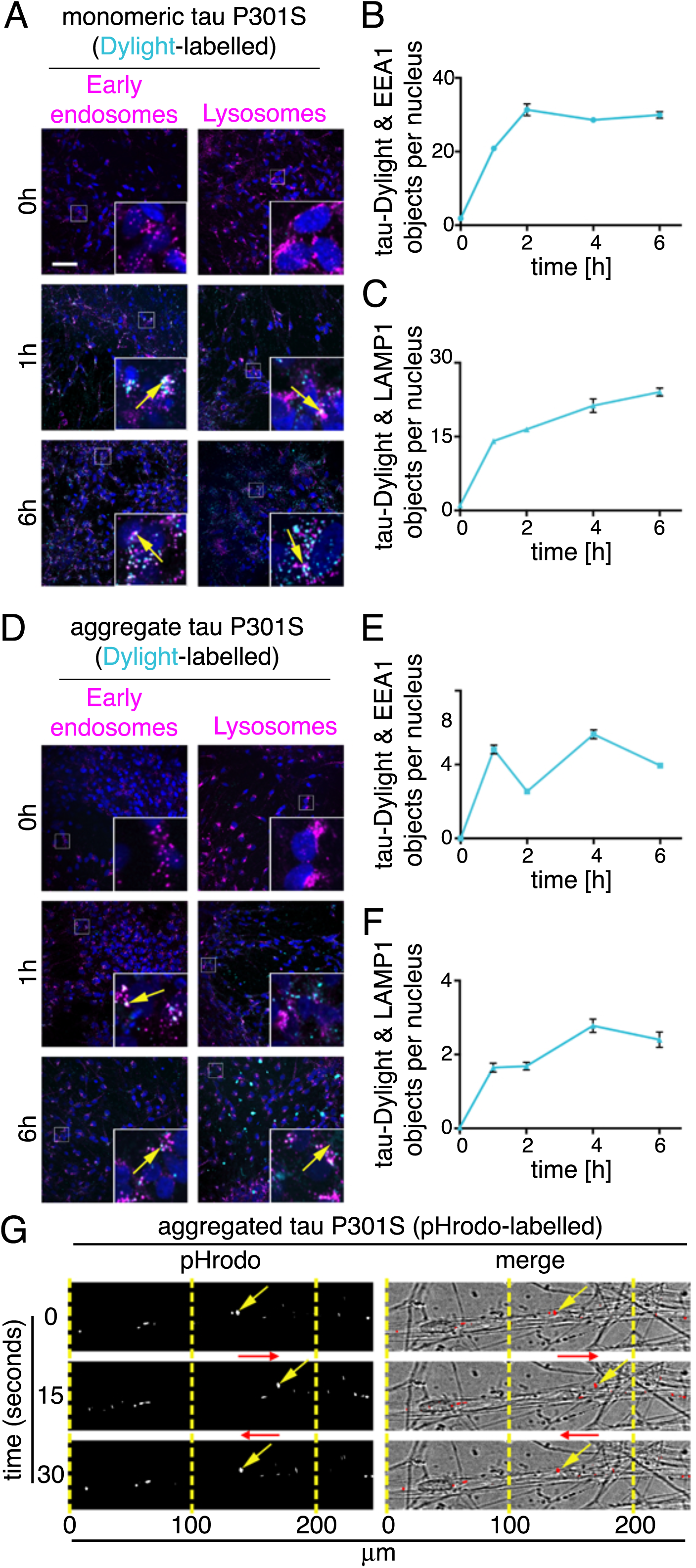
Monomeric and aggregated tau appear in early endosomes and lysosomes. Fixed immunofluorescent images of iPSC derived neurons (82 days after induction) after incubation with (A) 100nM monomeric tau P301S-Dylight (shown in cyan) for 0, 1 or 6 hours. Neurons washed with PBS were immunostained for EEA1 (early endosomal marker; left panel) or LAMP-1 (lysosome marker; right panel) and counterstained with nuclear marker (DAPI, shown in blue). Co-localised tau-Dylight and vesicular markers appears white in images and are indicated by arrows. Automated imaging was performed using an Opera Phenix platform; insets show magnifications of the area indicated in the main image. Scale bar, 50μm. Automated quantification (CellProfiler) of tau-Dylight and (B) EEA1 or (C) LAMP1 double positive objects per nucleus over time, 25 independent z-stacks were analysed per condition and time point, Error bars indicate standard deviation. (D-F) Analysis of co-localisation of 4.8 *μ*g.ml^−1^ (~100nM equivalent to monomeric) aggregated tau-Dylight with vesicular compartments over 6 hours, using the experimental conditions and parameters as (A-C). (F) Live tau-pHrodo imaging demonstrates the movement of vesicular structures in neurons. Representative images of neurons (80 days after induction) after 4 hours incubation with 1.2*μ*g.ml^−1^ of aggregated tau-pHrodo imaged at 1-second intervals, yellow arrows indicate intracellular dynamic tau-pHrodo positive objects, red arrows indicate the direction of movement.

In agreement with the monomeric tau-pHrodo experiments, monomeric tau-Dylight is rapidly taken up into neurons (Figure 3C). In contrast, IgG-Dylight shows little detectable entry into neurons throughout the time course (data not shown). As early as 1 hour after addition of extracellular tau, monomeric and aggregated tau-Dylight were co-localised in EEA1 early endosomes. Monomeric and aggregated tau-Dylight were also detected in LAMP1 late endosomes and lysosomes, consistent with endocytosed proteins first reaching the early endosomes, before endosomes mature to become late endosomes.

Live imaging (Figure 3G, Supplementary Movie 2) of neurons incubated with aggregated tau-pHrodo (four hours; 0.7*μ*g.ml^−1^) enabled the longer-term tracking of tau-pHrodo-containing vesicles within neurons. Tau-containing vesicles were dynamically and rapidly (>10*μ*m/second) transported in both antero- and retrograde directions along neurites, and could rapidly reverse direction of transport (Figure 3G). Thus, monomeric and aggregated tau both efficiently enter neurons via the endosome-lysosome system, and are actively trafficked within vesicles over long distances within neurons over several hours.

### Differential effects of dynamin inhibition on endocytosis of monomeric and aggregated tau

As extracellular monomeric and aggregated tau utilise endosomal pathways to enter neurons, but display different kinetics of uptake, we examined whether the underlying internalisation mechanisms of tau protein species differ. First, we tested the sensitively of monomeric and aggregated tau internalisation to the small molecule inhibitor Dynasore (Figure 4), an inhibitor of the GTPase Dynamin, which is required for numerous membrane fission events, including clathrin-mediated endocytosis (Preta et al., 2015). Following 30 minute pre-incubation with 100*μ*M Dynasore or vehicle control, entry of extracellular monomeric (25nM) or aggregated tau-pHrodo (1.2*μ*g.ml^−1^) was assessed by live imaging over four hours.

**Figure 4.**
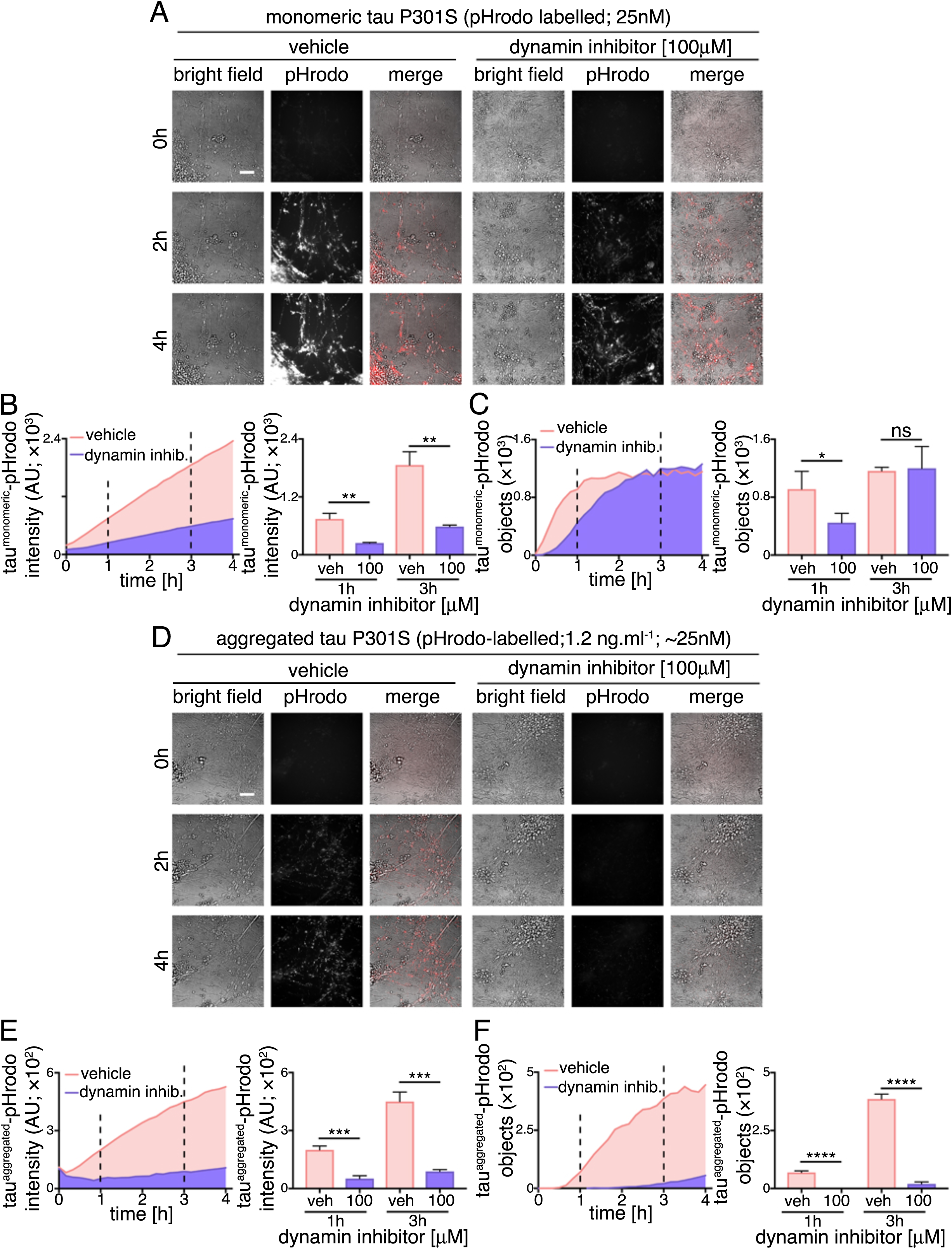
Differential effects dynamin inhibition on endocytosis of monomeric and aggregated tau. (A) Time-lapse (0-4 hours) images assessing dynamin inhibition on human neuronal internalization and acidification of extracellular monomeric and aggregated tau P301S-pHrodo. Neurons (75 days after induction) were preincubated with culture medium containing 100*μ*M Dynasore (dynamin inhibitior) or vehicle control (1% [v/v] DMSO) for 30 minutes, prior to exchanging culture medium to contain 1.2*μ*g.ml^−1^ extracellular tau-pHrodo and Dynasore or vehicle. Bright field (grey scale in merge) and pH sensitive fluorescent signal (pHrodo; red in merge) were captured using automated imaging on the Opera Phenix platform (Perkin Elmer). Nine independent measurements were taken from three technical replicates at 10 minutes intervals. Scale bar 100*μ*m. (B) Sum intensity of the pHrodo positive objects and (C) number of pHrodo positive objects was measured over 4 hours. Intensity and object measurements are displayed over  time and at a one- and three-hour time points (dashed lines). Analysis was performed using Harmony software (Perkin Elmer) error bars indicate standard deviation and and significance was determined using unpaired t test. (D-F) Effect of dynamin inhibition on internalization and acidification of extracellular aggregated tau-pHrodo into intraneuronal compartments using the same experimental conditions and parameters as (A-C).

The amount of monomeric tau entering neurons, as measured by total fluorescent intensity of intracellular monomeric tau-pHrodo vesicles, is significantly reduced by dynamin inhibition, as shown at 1 and 3 hours after addition of extracellular tau (Figure 4B). This is also reflected in the significant reduction in the number of tau-pHrodo positive objects at 1 hour (compared with vehicle control; Figure 4C). The kinetics of appearance of intracellular monomeric tau-pHrodo objects changes in the presence of dynamin inhibitor, displaying an initial lag in entry. However, the number of tau-pHrodo objects then reaches the same level as vehicle-treated controls by 3 hours, although not the same total intensity (Figure 4C), which may reflect a reduction in endosome/lysosome acidification in the presence of Dynasore. This suggests that there are two distinct mechanisms of entry of monomeric tau into human neurons, one of which is rapid and dynamin-dependent, and the other slower and independent of dynamin.

The effect of dynamin inhibition on entry of aggregated tau is more pronounced than on monomeric tau (Figure 4D-F). The total fluorescent intensity of intracellular aggregated tau-pHrodo is consistently reduced by more than 70% at 1 and 3 hours after tau addition (Figure 4E), and the number of tau-pHrodo positive objects is reduced by 95% (Figure 4F). These data indicate that, at the concentrations studied, that aggregated tau enters neurons almost entirely via clathrin-mediated endocytosis.

### Perturbation of actin dynamics confirms that monomeric and aggregated tau have overlapping but different routes to enter neurons

To explore further whether monomeric and aggregated tau enter neurons via different mechanisms, we studied the role of actin polymerisation in this process (Figure 5). Inhibition of actin polymerisation with Cytochalasin D disrupts several clathrin-independent endocytic processes, including bulk endocytosis/macropinocytosis (Mooren et al., 2012). Disruption of actin polymerisation has previously been shown to inhibit entry of fibrils formed of the tau repeat domain (Holmes et al., 2013). Therefore, neuronal entry of tau-pHrodo was measured over three hours in the presence of extracellular monomeric (25nM) or aggregated (1.2*μ*g.ml^−1^) tau-pHrodo, following 30 minute pre-incubation with 0.1 or 1*μ*M Cytochalasin D or vehicle control.

**Figure 5.**
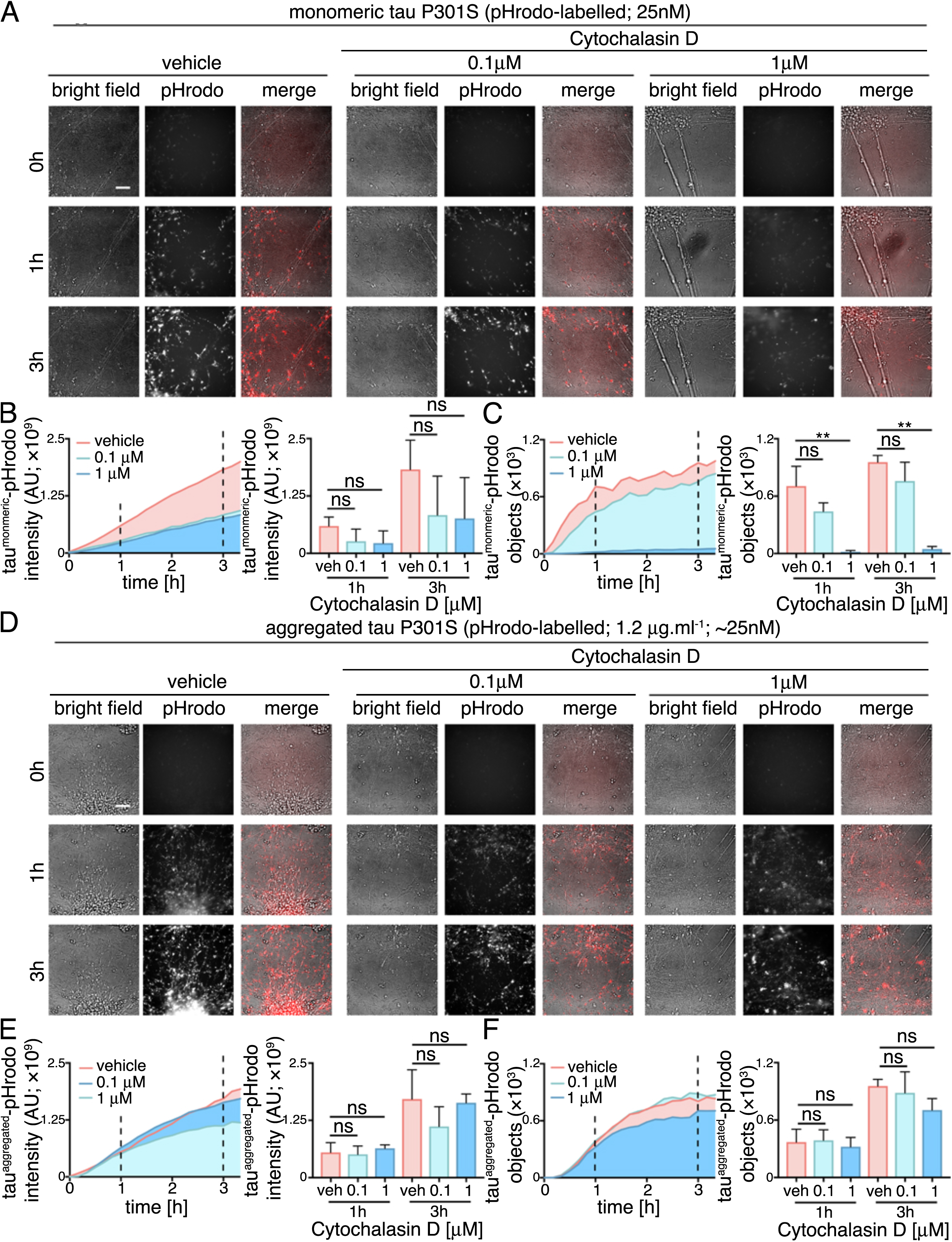
Perturbation of actin dynamics confirms that monomeric and aggregated tau enter neurons via different uptake mechanisms. (A) Time-lapse (0-3 hours) images assessing inhibition of actin polymerisation on human neuronal internalization and acidification of extracellular monomeric and aggregated tau P301S-pHrodo. Neurons (65 days after induction) were pre-incubated with culture medium containing 0.1, 1*μ*M Cytochalasin D or vehicle control (1% [v/v] DMSO) for 30 minutes, prior to exchanging culture medium to contain 1.2*μ*g.ml^−1^ extracellular tau-pHrodo and Cytochalasin D or vehicle. Bright field (grey scale in merge) and pH sensitive fluorescent signal (pHrodo; red in merge) were captured using automated imaging on the Opera Phenix platform (Perkin Elmer). Eight independent measurements were taken from three technical replicates at 10 minutes intervals. Scale bar 100*μ*m. (B) Sum intensity of the pHrodo positive objects and (C) number of pHrodo positive objects was measured over 3.3 hours. Intensity and object measurements are displayed over time and at a one- and three-hour time points (dashed lines). Analysis was performed using Harmony software (Perkin Elmer) error bars indicate standard deviation and significance was determined using one-way ANOVA. (D-F) Effect of inhibition of actin polymerisation on internalization and acidification of extracellular aggregated tau-pHrodo into human neurons using the same experimental conditions and parameters as (A-C).

Entry of monomeric tau was reduced in the presence of 1*μ*M Cytochalasin D, as reflected in the 95% reduction in the number of monomeric tau-pHrodo positive objects after three hours incubation in the presence of 1*μ*M Cytochalasin D (Figure 5C). In contrast, disruption of actin polymerisation with Cytochalasin D had little effect on entry of aggregated tau (total fluorescent intensity and number of objects; Figure 4D-F).

Live imaging of tau entry with pHrodo-tau may not detect intracellular pools of tau that enter via a different mechanism, avoiding low-pH environments. Additionally the pharmacological treatments may also effect the acidification of intracellular vesicles, which could affect the pHrodo signal. To control for this, we performed tau-Dylight internalisation assays in the presence of 100*μ*M Dynasore or 1*μ*M Cytochalasin D (Supplementary Figure 2), with three hours of incubation with 100*μ*M extracellular monomeric or aggregated tau. Compared with the live imaging studies, relatively high concentrations of aggregated tau were used to investigate whether the dynamin-dependence and actin-independence of neuronal entry seen by live imaging were related to the lower molarity of aggregated versus monomeric tau used for those experiments. These independent assays confirmed the same differential effects of the two inhibitors observed by live imaging of pHrodo-tau: at the three hour assay point, dynamin inhibition had no effect on the number of monomeric tau-Dylight positive punctae within neurons, whereas inhibition of actin polymerisation reduced the amount of intracellular tau by over half (Supplementary Figure 2). Conversely, dynamin inhibition significantly reduced the entry of aggregated tau, with no significant effects of Cytochalasin D (Supplementary Figure 2) at this relatively high molar concentration of aggregated tau.

### Anti-tau antibodies slow neuronal internalisation of extracellular tau

It is not currently known if tau entry into neurons requires specific domains of tau, specific cell surface binding proteins or receptors. Antibodies that recognise extracellular tau are a potential therapeutic strategy for slowing disease progression through the CNS by inhibiting tau entry into neurons (Congdon et al., 2016; Gu and Sigurdsson, 2011). Complex formation between specific antibodies and tau results in an increase in molecular size and may alter or mask uptake recognition sites on tau, thus preventing neuronal entry of tau.

We analysed the ability of a polyclonal antibody to the C-terminal half of tau to reduce tau protein entry into human cortical neurons. The anti-tau antibody or control IgGs (250nM) were incubated with either monomeric (25nM) or aggregated (1.2*μ*g.ml^−1^) tau-pHrodo at approximately 10-fold excess of antibody for 30 minutes. Tau-antibody mixtures were then added to iPSC-derived human neurons and tau entry assessed by live imaging of fluorescent intracellular tau-pHrodo (Figure 6).

**Figure 6.**
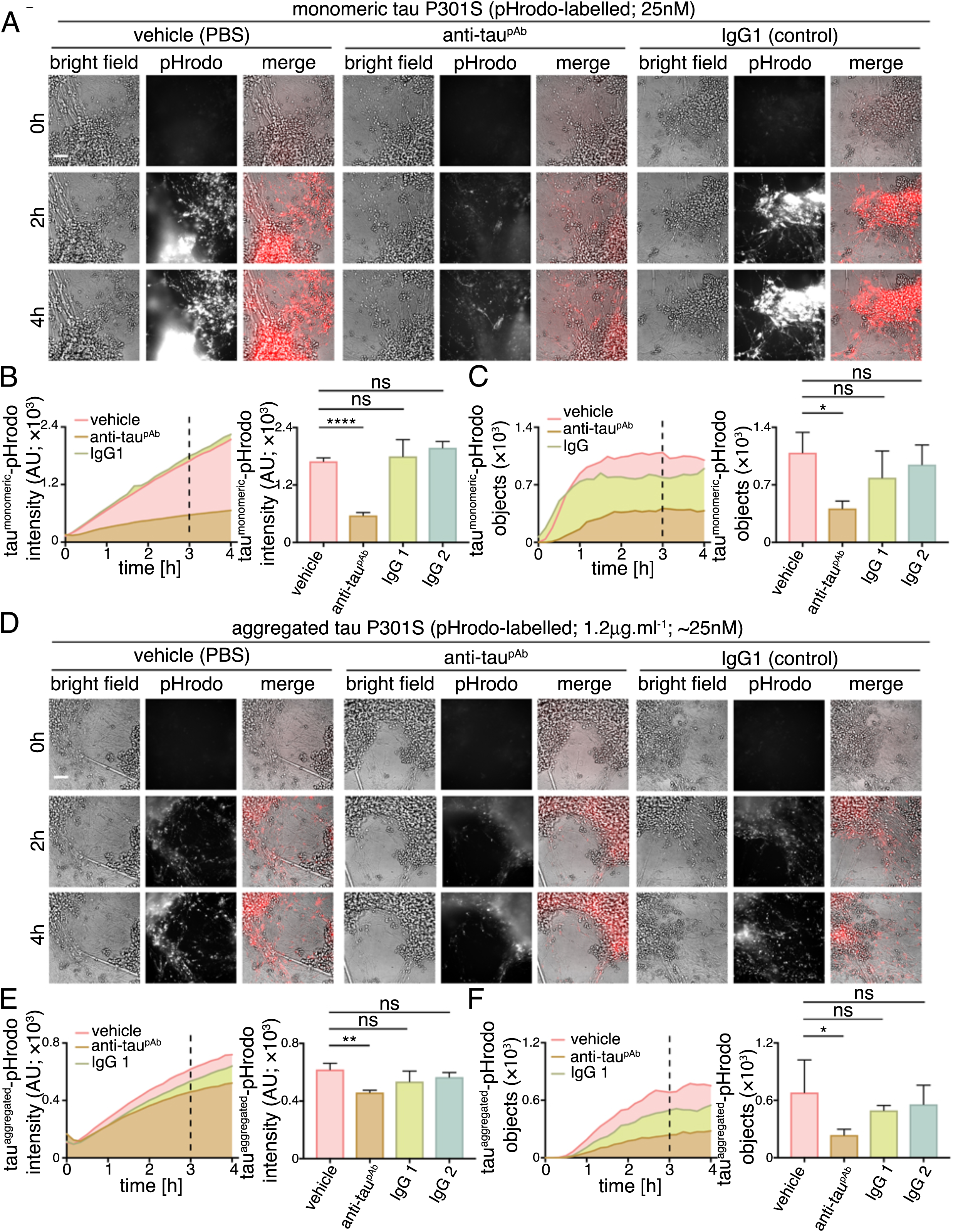
Anti-tau antibodies have differing effects in slowing neuronal uptake of extracellular tau. (A-C) Time-lapse (0-4 hours) imaging to monitor the effect of anti-tau polyclonal antibody on internalization and acidification of extracellular monomeric tau-pHrodo by human neurons. Monomeric tau-pHrodo (25nM) was pre-incubated with culture medium containing 250nM of polyclonal antitau (anti-tau^pAb^), IgG control (IgG1) antibodies or vehicle control (PBS) for 30 minutes. Tau-antibody incubations were added to iPSC-derived human neuronal cultures (75 days after induction). Bright field (grey scale in merge) and pH sensitive fluorescent signal (pHrodo; red in merge) were captured using automated imaging on the Opera Phenix platform (Perkin Elmer). Eight independent measurements were taken from three technical replicates at 10 minutes intervals. Scale bar 100*μ*m. (B, C) Intensity and number of pHrodo-positive objects measured over four hours after addition of extracellular monomeric tau-pHrodo in the presence of vehicle, anti-tau polyclonal antibody or two different control IgG antibodies. Intensity and object measurements are displayed over time and at a three-hour time point (dashed line). Analysis was performed using Harmony software (Perkin Elmer) error bars indicate standard deviation and and significance was determined using one-way ANOVA. (D-F) Effect of tau antibody on internalization and acidification of extracellular aggregated tau-pHrodo into human neurons using the same experimental conditions and analysis parameters as for monomeric tau in (AC).

Pre-incubation with the polyclonal (anti-tau^pAb^) tau antibody reduced the amount of monomeric tau that enters neurons, as assessed by the number of monomeric tau-pHrodo containing vesicles (Figure 6B; Supplementary Figure 3). The kinetics of monomeric tau entry are affected by the presence of tau-specific antibodies, such that an initial lag in the appearance of defined tau-pHrodo-positive objects was observed (Figure 6B).

Tau antibodies also reduce entry of aggregated tau uptake into human neurons (Figure 6C). In the presence of anti-tau^pAb^ the number of aggregated tau-pHrodo vesicles was reduced by more than half (Figure 6D; Supplementary Figure 3). IgG antibodies that do not recognise tau appear to have some minor effects on monomeric and aggregated tau entry, which may be due to competition between tau and IgG for bulk endocytosis.

The effect of tau antibodies on the entry of monomeric tau-Dylight into neurons was also tested (Supplementary Figure 3), using the same concentrations and pre-incubation time as the tau-pHrodo assays. After three hours of incubation with antibodies and extracellular monomeric or aggregated tau, neurons were washed, fixed and number of intracellular tau-Dylight punctae were quantified. These independent assays confirm that tau-specific antibodies significantly reduce the amount of tau-Dylight entering neurons (Supplementary Figure 3).

### Entry of aggregated tau-antibody complexes into neurons

Although tau was pre-incubated with a large molar excess of antibodies (10x) before addition to neurons, a notable amount of both monomeric and aggregated tau still entered neurons. This raised the question whether the tau that entered neurons under these conditions did so in a complex with antibodies, or that a fraction of antibody-free tau was available for neuronal entry. To distinguish between these possibilities, we performed additional assays using pre-formed aggregated tau-Dylight-antibody complexes. After three hours of incubation, antibodies were detected in the fixed neurons with a secondary antibody specific to the host species (anti-rabbit polyclonal; Figure 7) and also imaged for tau-Dylight. Intraneuronal punctae positive for both tau-Dylight and tau specific antibodies were detected after binding of extracellular tau with anti-tau^pAb^, demonstrating that tau-antibody complexes do enter human neurons. No intraneuronal tau-antibody complexes were detected when aggregated tau was incubated with IgG control.

**Figure 7.**
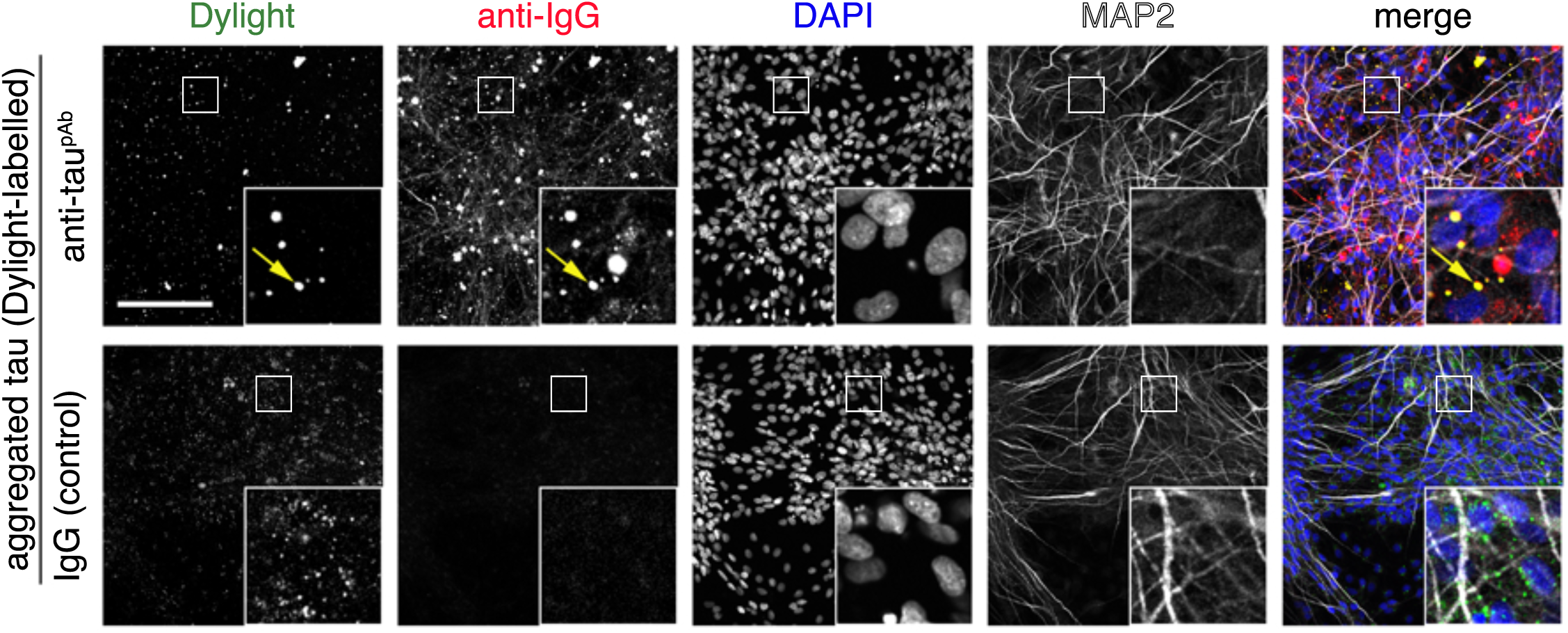
Aggregated tau endocytosed in the presence of antibody carries antibody into neurons. Immuno-fluorescent images monitoring neuronal uptake of extracellular aggregated tau P301S-Dylight-antibody complexes. Pre-formed (30 minutes incubation) aggregated tau-Dylight-antibody (anti-tau^pAb^ or an IgG control) complexes were added to iPSC derived neurons (71 days after induction). After three hours of incubation, neurons were washed, fixed and co-stained with antibody specific to the pre-incubation antibodies (donkey anti mouse IgG Alexa 594 or donkey anti rabbit IgG Alexa 594), nuclear (DAPI; blue in merge) and dendritic markers (MAP2; white in merge). Fluorescent signals were captured using using both Olympus FV1000 confocal and representative images are shown. Scale bar 100*μ*m.

## Discussion

We find here that both monomeric and aggregated tau (P301S) efficiently enter human neurons, arguing that tau entry to neurons is a constitutive biological process, and not a disease-specific phenomenon. Both species of tau enter neurons via a dynamin-dependent process, with monomeric tau also entering neurons through a second, slower pathway dependent on actin polymerisation. Monomeric and aggregated tau access neurons via the lysosome/endosome system, and a fraction of tau is dynamically transported long distances in neurons within low pH vesicles for lengthy periods. The amount of monomeric and aggregated tau entering neurons is reduced by polyclonal anti-tau antibodies. However, a notable amount of tau enters neurons complexed with antibodies, even in the presence of a ten-fold molar excess of antibodies. While antibody binding may affect intracellular trafficking of tau and the final destination of tau within the neuron (McEwan et al., 2017), these data demonstrate that simple binding of tau by antibodies is not sufficient to prevent neuronal entry when tau is presented in solution to the neuron.

In contrast to earlier reports from HEK cells and primary rodent neurons (Frost et al., 2009; Holmes et al., 2013; Wu et al., 2013), full-length monomeric tau efficiently and rapidly enters healthy human neurons, and does so as efficiently as aggregated tau. We find that there are two mechanisms of entry of monomeric tau to neurons: a rapid early phase that can be blocked by dynamin inhibition, and a second, slower phase that it does so in an actindependent manner. These two mechanisms suggest that monomeric tau enters neurons via a rapid saturable clathrin-mediated endocytic mechanism and also by bulk endocytosis (Loebrich, 2014). In contrast to monomeric tau, aggregated tau entry is independent of actin polymerisation and largely dynamin dependent, consistent with clathrin-mediated endocytosis. The lack of actin dependence of aggregated tau entry into neurons suggests that it does not enter human neurons via macropinocytosis, as has been reported in non-neuronal cells (Holmes et al., 2013).

Rapid clathrin-mediated endocytosis of both monomeric and aggregated tau would typically require one or more specific receptors or cell surface binding molecules, the identities of which are currently unknown. Specific surface receptors have been identified for fibrils of alpha-synuclein (Mao et al., 2016), and that receptor shows specificity for aggregated synuclein over the monomeric form, arguing that although both species of tau enter neurons via clathrin-mediated endocytosis, they may do so through distinct surface receptors. As a largely disordered protein, the most obvious candidate region for recognition for cellular entry of monomeric tau is the highly ordered microtubule-binding region of tau (Butner and Kirschner, 1991). Detailed understanding of the different paths of entry for monomeric and aggregated tau will be useful for investigating the potential for specifically preventing inter-neuronal transfer of aggregated tau without interfering with transfer of non-pathogenic forms.

In addition to the importance of these questions for understanding mechanisms of dementia pathogenesis, they also have implications for immunotherapy strategies that target extracellular tau as disease-modifying treatments for dementia. Tau immunotherapy strategies have been shown to alter tau transmission in mouse models (Congdon et al., 2016; Gu and Sigurdsson, 2011). Proposed mechanisms for tau immunotherapies are epitope, affinity and aggregate size-dependent. Studies of specific tau immunotherapies have demonstrated that either extracellular (Yanamandra et al., 2013) or intracellular (Gu et al., 2013; Nicholls et al., 2017) tau can be targeted. Antibodies are thought to halt the progression of disease by binding to tau aggregates, triggering their uptake and clearance via either endosomal or proteasome pathways (Chai et al., 2011; Gu et al., 2013; Mallery et al., 2010). Fluid phase (Funk et al., 2015) and Fc-receptor-mediated (Congdon et al., 2016) endocytosis have both been implicated in the uptake of antibody-tau complexes into neurons and microglia. The consequences for the neuron of uptake of antibody-antigen complexes are not clear, and would depend in part on whether those complexes are degraded or accumulate in the endosome/lysosome or transit to the cytoplasm, where they could be detected by the cytosolic Fc receptor TRIM21 and targeted for degradation (McEwan et al., 2017).

The data presented here from in vitro experiments suggest that entry of tau into human neurons is an efficient physiological process, rather than a disease-specific gain-of-function. Most current immunotherapy approaches targeting extracellular tau do not distinguish between pathogenic forms of tau that are thought to propagate disease, and the forms of extracellular tau that are found in the healthy brain (Bright et al., 2015). Disruption of interneuronal transfer of non-pathogenic tau may have deleterious effects, if that transfer has a biological function. Current clinical trials of anti-tau antibodies have not so far reported deleterious effects relating to disruption of interneuronal non-pathogenic tau, and the field awaits the outcomes of longer term trials. Furthermore, therapeutics that are not selective for extracellular pathogenic forms of tau may not achieve appropriate levels of target engagement in the presence of concentrations of non-pathogenic tau that are higher than those of pathogenic tau. Therefore, it will be important to ascertain the biological functions of neuronal release and internalisation of tau, and to identify which, if any, extracellular tau forms are truly pathogenic, to facilitate the rational design of therapies that target interneuronal transfer of tau to prevent disease progression.

## ACKNOWLEDGEMENTS

Research in the FJL group at the Gurdon Institute is supported by the Alborada Trust (ARUK Stem Cell Research Centre) and Dementias Platform UK. FJL is a Wellcome Trust Investigator.

## AUTHOR CONTRIBUTIONS

Experimental work was performed by TW, LE, GF and JS, with data analysis by TW, LE and FJL. The study was designed by LE, TW, FJL, MP and AB. All authors contributed to the manuscript writing.

## Methods

### Production and characterisation of human iPSC-derived cerebral cortex neurons

Directed differentiation of hESCs and iPSCs to cerebral cortex was carried out as described, with minor modifications (Shi et al., 2012a; Shi et al., 2012b). For drug treatment, all compounds were dissolved in DMSO at the concentrations noted, and DMSO was the vehicle control in all experiments. Compounds were added 30 minutes prior to incubation with recombinant tau protein: Dynamin inhibitor, Dynasore (abcam); Actin polymerisation inhibitor, Cytochalasin D (Tocris Bioscience).

### Recombinant tau expression and purification

Tau P301S_10Xhis-tag_avi-tag was over expressed in BL21(DE3) bacteria. Cells were lysed using BugBuster (Millipore) and clarified lysate was applied to a 5ml HisTrapHP column (GE Healthcare) in 2×PBS. Tau was eluted using a 0-500mM imidazole gradient. Peak fractions were pooled and further purified in 2×PBS using a Superdex 200 16/60 gel filtration column (GE Healthcare). Pooled factions were then concentrated to approximately 8mg/ml using a spin concentrator (Millipore). Final protein concentration was determined by Nanodrop analysis.

### Preparation of tau oligomeric species

One millilitre of tau P301S at 8mg/ml was incubated with 4mg.ml^−1^ heparin (Sigma) in PBS plus 30mM MOPS pH7.2 at 37^o^C for 72 hours. Aggregated material was diluted in 9ml of PBS plus 1% (v/v) sarkosyl (Sigma) and left rocking for 1 hour at room temperature to completely solubilize any non-aggregated material. Insoluble tau was pelleted by ultracentrifugation at 50,000 rpm in a TLA70.1 rotor for 1 hour at 4^o^C. The pellet was resuspended in 1ml of PBS by vigorous pipetting and sonicated at 100W for 3×20 seconds using a Hielscher UP200St ultrasonicator to disperse clumps of protein and break large filaments into smaller species.

### Labelling of purified recombinant tau

To label purified recombinant tau with Dylight 488 NHS ester (Thermo Fisher), monomeric or aggregated tau P301S (concentration of 8 and 2 mg.ml^−1^, respectively) was dialysed into PBS using Slide-A-Lyzer mini dialysis units (Thermo Fisher) overnight at 4°C. The dialysed tau preparation was then conjugated to 50*μ*g of Dylight according to the manufacturer’s protocol. Non-incorporated dye was removed by overnight dialysis against 2×11 of PBS at 4°C and the concentration determined using a BCA protein quantification kit (Thermo Fisher).

To label purified recombinant tau with a pH-sensitive form of rhodamine (pHrodo; Thermo Fisher Scientific), 150*μ*M of tau protein (or equivalent protein concentration for aggregate; ~7*μ*g.ml-1) was incubated with 1.5mM of pHrodo Red Maleimide (dissolved in DMSO) and 1.5mM of TCEP (1:10:10 molar ratio, respectively) for 2hrs in the dark at RT. After incubation, labelled protein samples were subjected to size exclusion chromatography at 4°C (Superdex 200 Increase 10/300 GL; GE healthcare) in 50mM phosphate (pH7.4), 150mM NaCl, to remove unreacted dye and assess perturbation in oligomeric state by labelling (no change was observed, data not shown). 10mM 10-kDa dextran-conjugated to pHrodo (Thermo Fisher Scientific) was dissolved in DMSO.

### Labelled tau internalization assays

Tau P301S labelled with either Dylight or pHrodo was diluted in neuronal maintenance medium tissue culture medium and added to neurons at the concentration and for the time indicated. Neurons were either washed three times using 1xHBSS before processing for immunofluorescence, processing for flow cytometry or subjected to live imaging.

### Immunostaining, confocal microscopy and data analysis

Immunohistochemistry was performed on neurons as previously described (Shi et al., 2012) using the following primary antibodies: MAP2 from Abcam (ab5392); EEA1 and LAMP1 from Cell Signaling (C45B10 and D2D11). Secondary antibodies used were goat anti mouse Alexa594 (A21125), goat anti chicken Alexa 647 (A21449), goat anti rabbit Alexa 546 (A11010), donkey anti mouse Alexa 594 (A21203), donkey anti rabbit Alexa 594 (A21207, all from Thermo Fisher). Images were acquired using Olympus FV1000 or Opera Phenix. Typically conditions were repeated in triplicate and >12 images were collected per well. Quantification of fixed cell imaging was performed using CellProfiler.

### Flow Cytometry

After three 1xHBSS washes neurons were dissociated using Accutase solution (Sigma, A6964) into single cells. Dissociated cells were collected by low speed centrifugation (1000xg for 1min), the supernatant removed and cells were fixed using 4% formaldehyde in PBS for 20min at room temperature. Cells were washed 3x with PBS, nuclei counterstained with DAPI and analysed using a BD LSRFortessa. Flow cytometry data was analysed using ‘Flowing’ software.

### Live imaging and data analysis

Neurons grown on μ-Plate 96 well (Ibidi) were imaged for fluorescent pHrodo labelled protein (excitation at 561nm, emission at 570-630nm) in an acidic environment using a 40× confocal Opera Phenix high content screening system (Perkin Elmer). Bright field and fluorescence emission images were collected at 10-minute intervals. Parameters for fluorescent objects were set (fluorescent intensity and contrast) and quantified using the Harmony software (Perkin Elmer). Data were analysed using Prism Software (GraphPad). Typically, conditions were repeated at least in triplicate and <4 (typically 8) fields were recorded per well. To track individual objects images were collected at 1-second intervals.

